## Supplementary Information for "An ancestral dual function of OmpM as outer membrane tether and nutrient uptake channel in diderm Firmicutes"

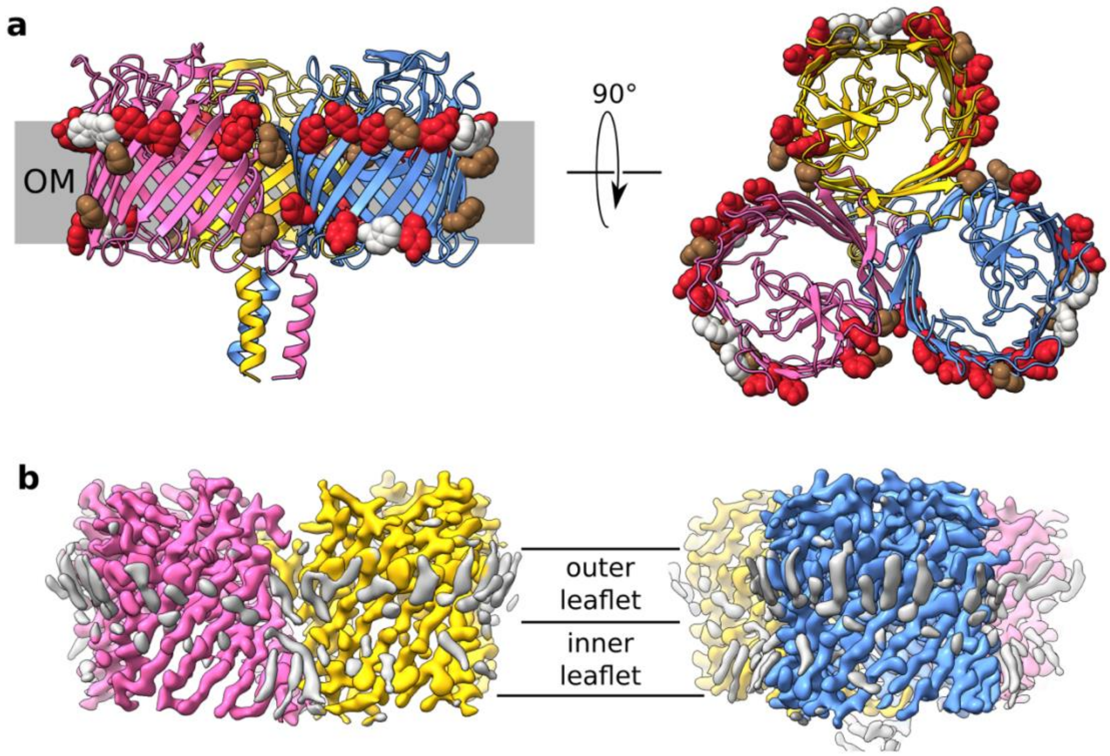


**Supplementary Figure 1. VpOmpM1 aromatic girdle and putative lipid density. a** Aromatic residues forming the aromatic girdle shown in space-filling representation on the cartoon of C1 VpOmpM1 reconstruction. Brown, phenylalanine; red, tyrosine; grey, tryptophan. **b** Cryo-EM density of the C3 VpOmpM1 reconstruction. Protein density coloured by chain; grey, putative lipid or detergent density.

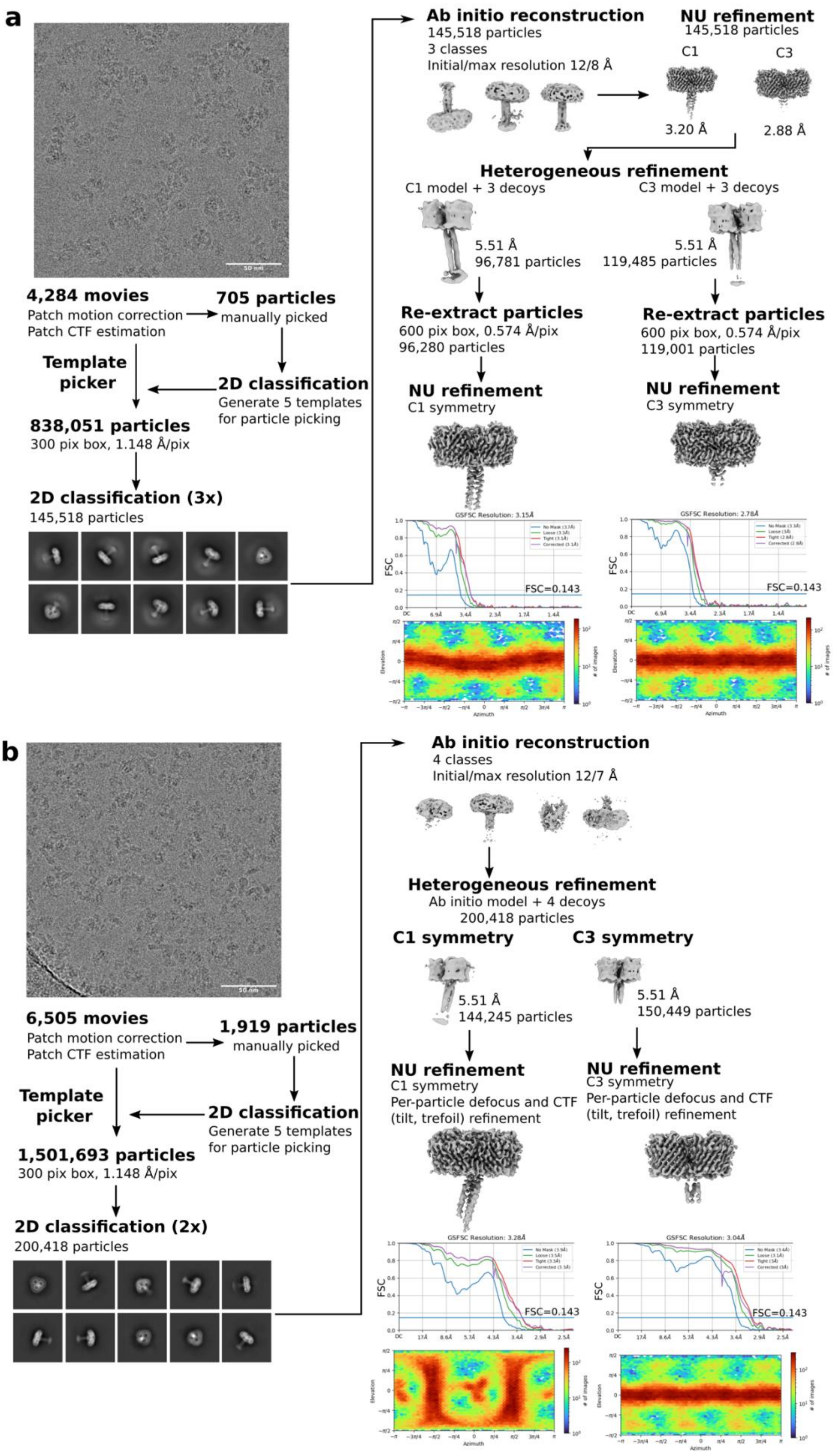

a
4,284 movies
Patch motion correction
Patch CTF estimation
705 particles
manually picked
Template picker
2D classification
Generate 5 templates for particle picking
838,051 particles
300 pix box, 1.148 Å/pix
2D classification (3x)
145,518 particles
Ab initio reconstruction
145,518 particles
3 classes
Initial/max resolution 12/8 Å
NU refinement
145,518 particles
C1
C3
3.20 Å
2.88 Å
Heterogeneous refinement
C1 model + 3 decoys
C3 model + 3 decoys
5.51 Å
96,781 particles
5.51 Å
119,485 particles
Re-extract particles
600 pix box, 0.574 Å/pix
96,280 particles
Re-extract particles
600 pix box, 0.574 Å/pix
119,001 particles
NU refinement
C1 symmetry
NU refinement
C3 symmetry
GSFSC Resolution: 3.15Å
GSFSC Resolution: 2.78Å
FSC
FSC=0.143
Elevation
Azimuth
### of images
b
6,505 movies
Patch motion correction
Patch CTF estimation
1,919 particles
manually picked
Template picker
2D classification
Generate 5 templates for particle picking
1,501,693 particles
300 pix box, 1.148 Å/pix
2D classification (2x)
200,418 particles
Ab initio reconstruction
4 classes
Initial/max resolution 12/7 Å
Heterogeneous refinement
Ab initio model + 4 decoys
200,418 particles
C1 symmetry
C3 symmetry
5.51 Å
144,245 particles
5.51 Å
150,449 particles
NU refinement
C1 symmetry
Per-particle defocus and CTF (tilt, trefoil) refinement
NU refinement
C3 symmetry
Per-particle defocus and CTF (tilt, trefoil) refinement
GSFSC Resolution: 3.28Å
GSFSC Resolution: 3.04Å
FSC
FSC=0.143
Elevation
Azimuth
### of images

**Supplementary Figure 2. Cryo-EM data processing**. Cryo-EM data processing workflows for VpOmpM1 expressed in *E. coli* (**a**) and purified from *V. parvula* (**b**), showing representative motion-corrected movies and 2D class averages; intermediate and final cryo-EM maps; Fourier shell correlation (FSC) curves and particle angular distribution (elevation vs azimuth) plots for the final maps.

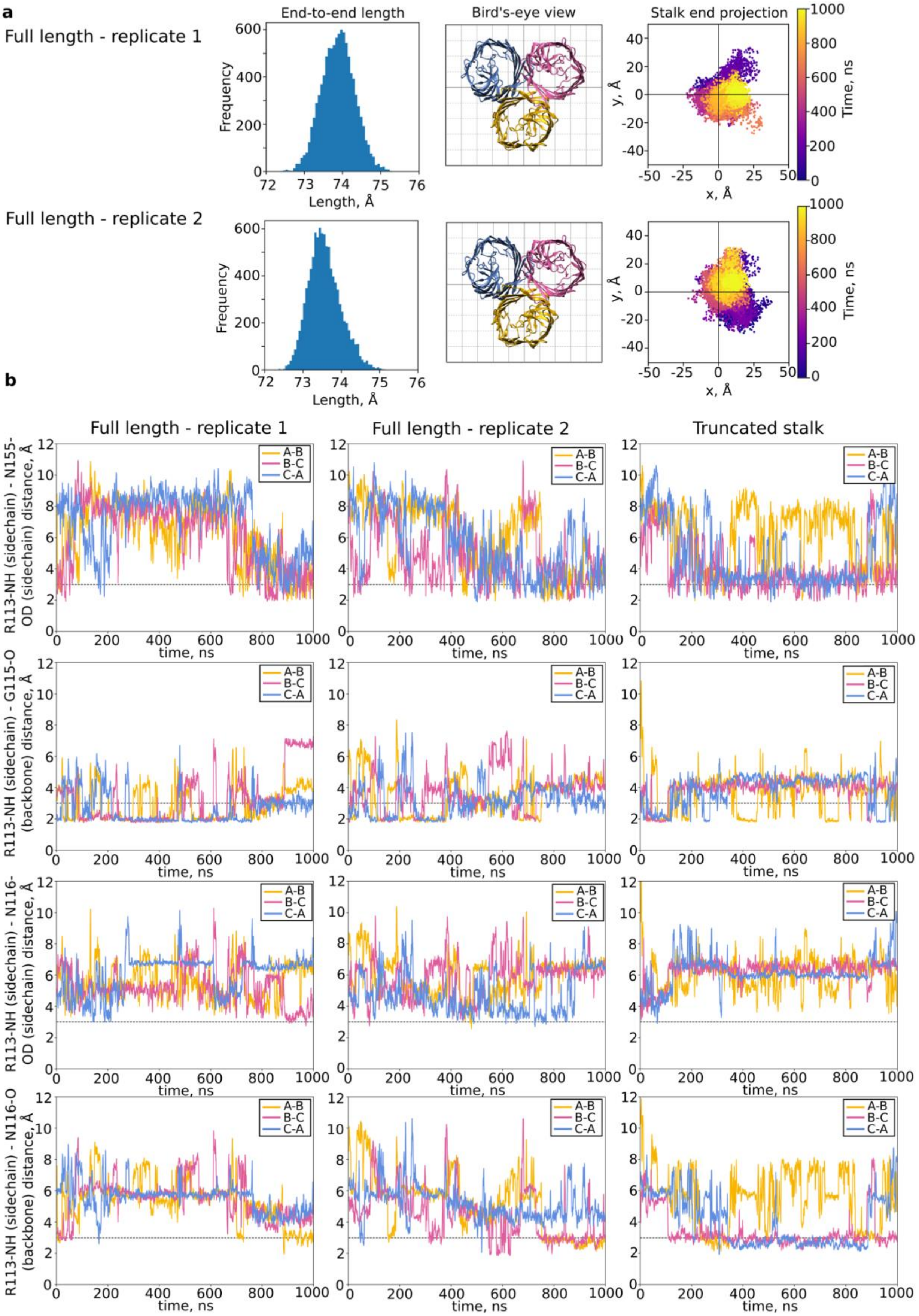


**Supplementary Figure 3. All-atom molecular dynamics simulation replicates**. **a** Stalk length and end projection plots for the two replicates of all-atom simulations with the native VpOmpM1 and AlphaFold2 graft model (full-length). The bird's-eye-view coordinates correspond to the plot coordinates in the stalk projection plot. **b** Hydrogen bond distances throughout the simulation. The ‘truncated stalk’ simulation was performed with the native VpOmpM1 model without the AlphaFold2 graft. Data for replicate 1 shown in this figure are the same as the data shown in Figure 2.

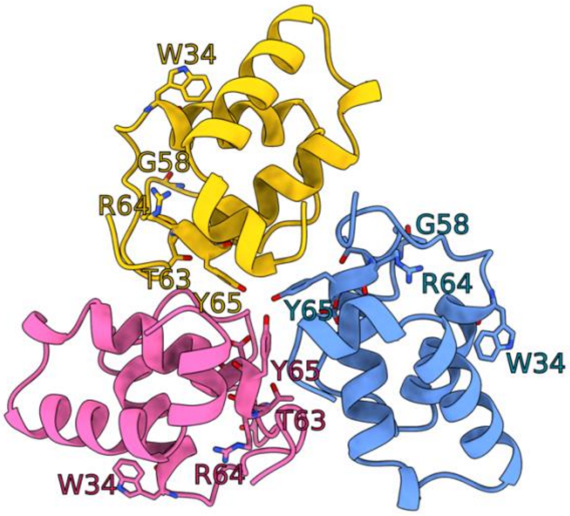


**Supplementary Figure 4. Putative PG-binding motifs in the SLH crystal structure**. Residues that are part of motifs conserved in SLH domains are shown in stick representation. Motif residues are not located in intra-protomer grooves as seen in crystal structures of SLH domains from Gram-positive Firmicutes and in the extended conformation of the VpOmpM1 stalk predicted by AlphaFold2 (Figure 3).

| Rank | Chain | Z | rmsd | lali | nres | %id | Description |
|---|---|---|---|---|---|---|---|
| 1 | 5f7l-C | 5.7 | 2.5 | 57 | 375 | 11 | Blood group antigen binding adhesin BabA |
| 2 | 6bml-B | 5.2 | 3.3 | 67 | 294 | 12 | Human palmitoyltransferase DHHC20 |
| 3 | 6fws-B | 5.2 | 3.3 | 68 | 683 | 4 | Helicase DinG |
| 4 | 6gmm-A | 5.2 | 2.9 | 61 | 433 | 11 | Adhesin LabA |
| 5 | 6fws-A | 5.0 | 2.7 | 65 | 686 | 8 | Helicase DinG |
| 6 | 7khm-A | 4.8 | 3.3 | 68 | 290 | 12 | Human palmitoyltransferase DHHC20 |
| 7 | 5f7w-A | 4.8 | 2.7 | 60 | 419 | 10 | Blood group antigen binding adhesin BabA |
| 8 | 5f7y-A | 4.8 | 3.2 | 62 | 417 | 10 | Blood group antigen binding adhesin BabA |
| 9 | 5has-A | 4.7 | 3.2 | 64 | 381 | 5 | DCB-HUS domain of Sec7 |
| 10 | 5f8q-A | 4.7 | 2.4 | 56 | 419 | 11 | Blood group antigen binding adhesin BabA |

**Supplementary Figure 5. SLH domain crystal structure DALI search results**. DALI [1] analysis shows that the VpOmpM1 SLH domain crystal structure has low similarity to other proteins from the PDB.

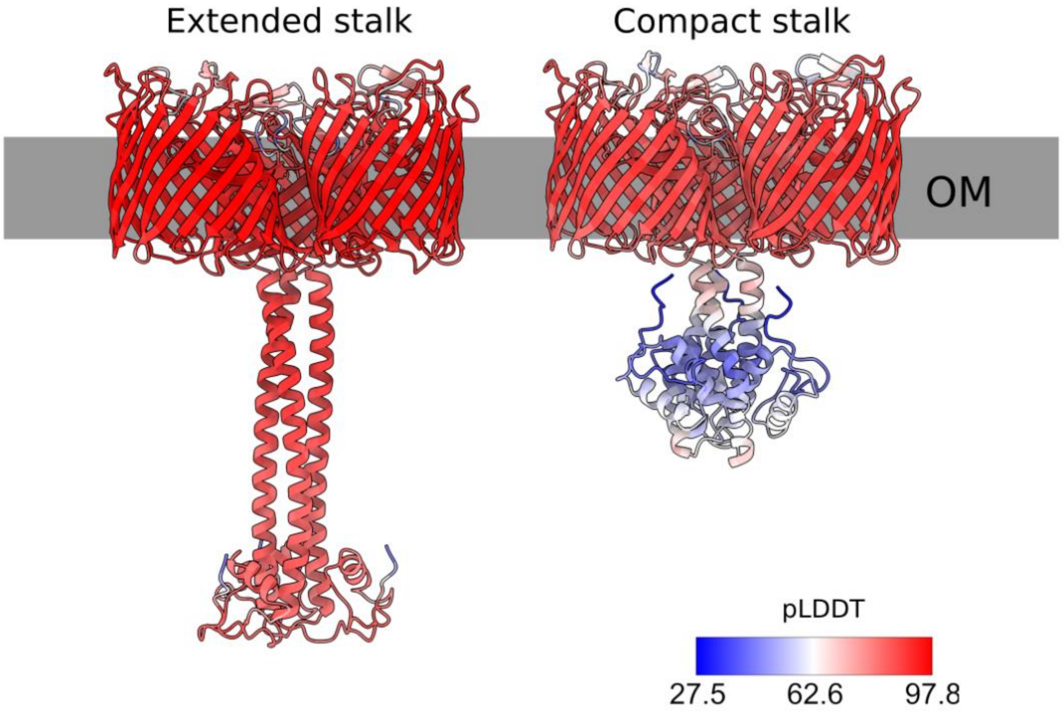


**Supplementary Figure 6. VpOmpM1 trimer AlphaFold2 [2] predictions**. Cartoons coloured by per residue confidence score (colour key): pLDDT>90 – very high; 90>pLDDT>70 confident; 70>pLDDT>50 low; pLDDT<50 – very low.

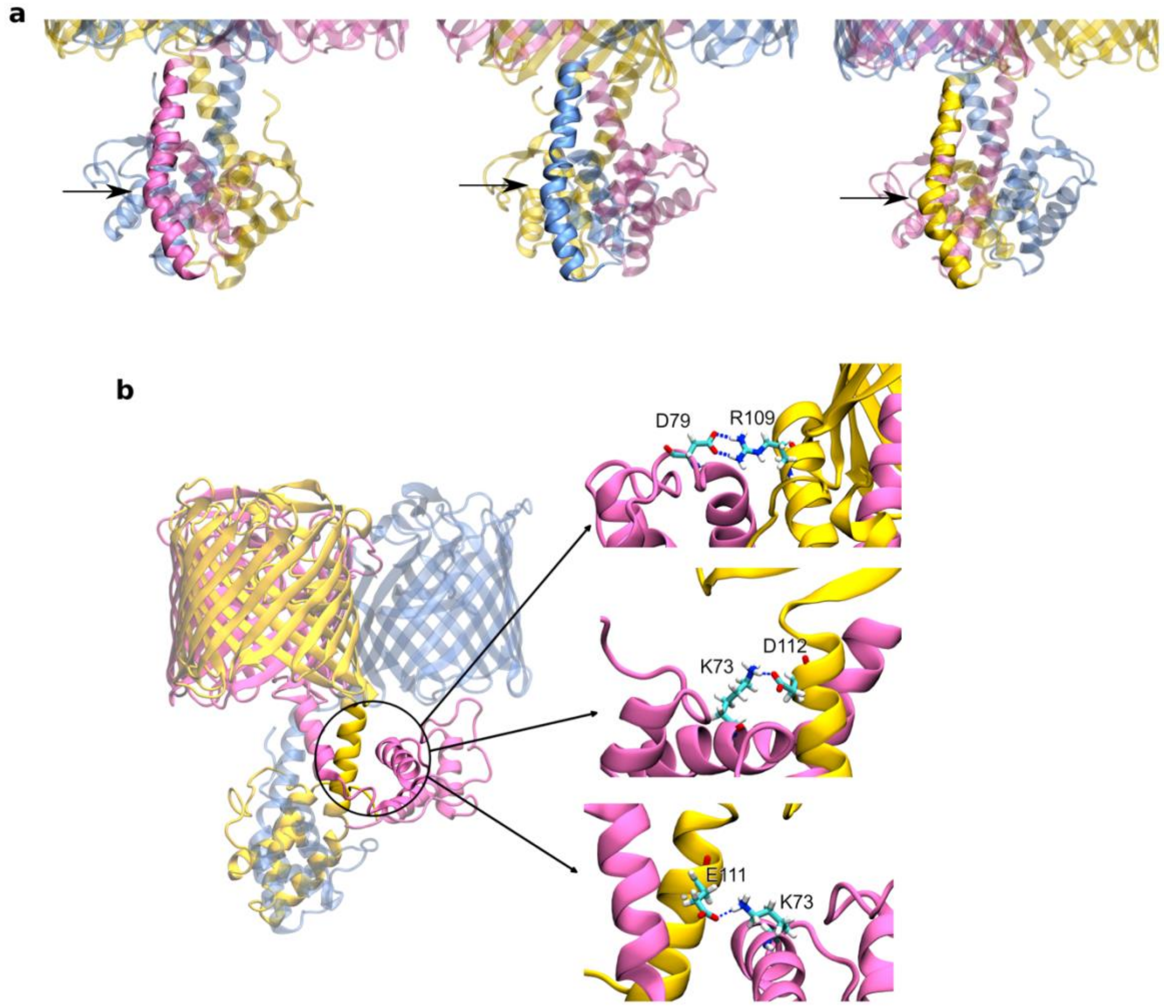


**Supplementary Figure 7. VpOmpM1 compact stalk simulation**. **a** The stalk $\alpha$–helices in each protomer are kinked at residues 97-102 (arrows). This kink is the hinge that allows movement of the stalk towards the OM, as seen for the pink protomer in the simulation. **b** The conformation of the pink protomer is stabilised by salt bridge formation with the stalk of the yellow protomer. Salt bridge occupancy throughout the simulation (top to bottom): 14.33%, 3.21% and 1.27%.

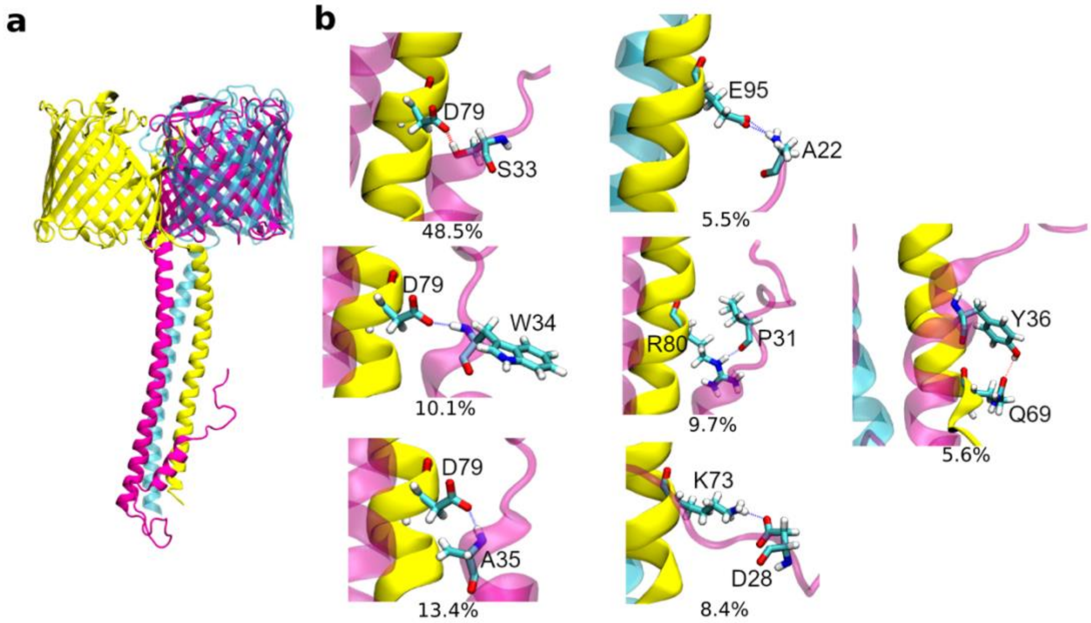


**Supplementary Figure 8. Interaction of SLH domain with coiled-coil during extended stalk MD simulations. a** During one all-atom simulation of VpOmpM1 with the grafted stalk, the SLH domain of one protomer (pink) unfolded and formed contacts with the coiled-coil region of another protomer (yellow). The SLH domains of the yellow and cyan protomers are not shown. **b** Close-up views of the SLH-coiled coil interactions. The occupancy of each interaction throughout the 1 μs simulation is shown under each panel.

**a**

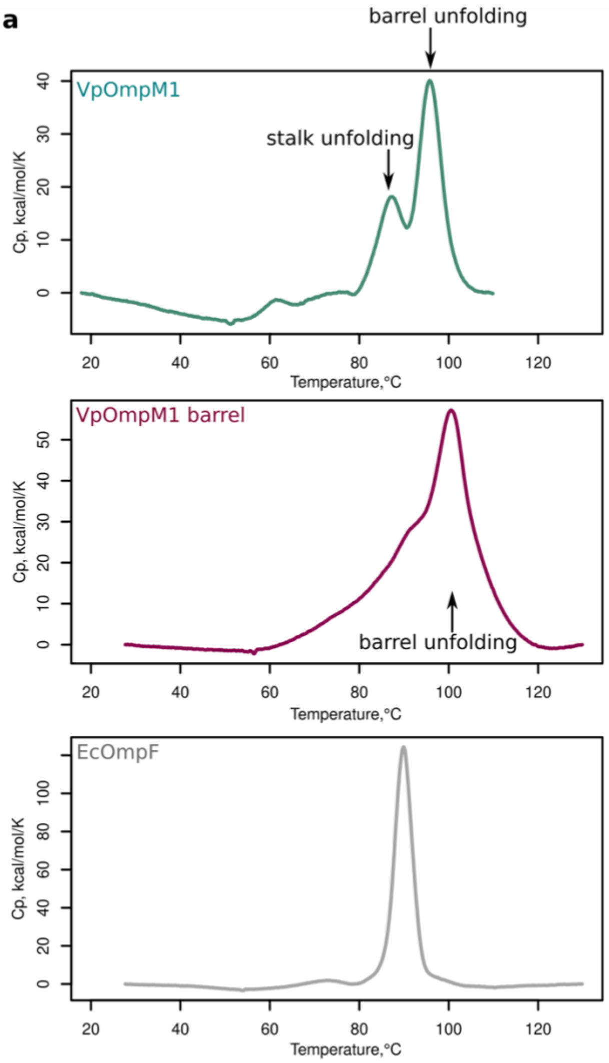


**b**

| Sample | Melting temperature | |
|---|---|---|
| | $T_m1$ | $T_m2$ |
| VpOmpM1 | 87.46°C | 95.74°C |
| VpOmpM1 barrel | 92.27°C | 101.06°C |
| EcOmpF | 89.93°C | N/A |

**Supplementary Figure 9. Protein melting temperature analysis**. **a** Dynamic scanning calorimetry thermograms for full-length VpOmpM1, VpOmpM1 barrel domain, and EcOmpF. Measurements were performed on a Malvern VP Capillary DSC instrument, using 18.7 μM protein in 10 mM HEPES-NaOH pH 7.0, 100 mM NaCl and 0.12% DM in each experiment. **b** Data were fitted and the melting temperatures ($T_m$) were extracted via the instrument manufacturer's software.

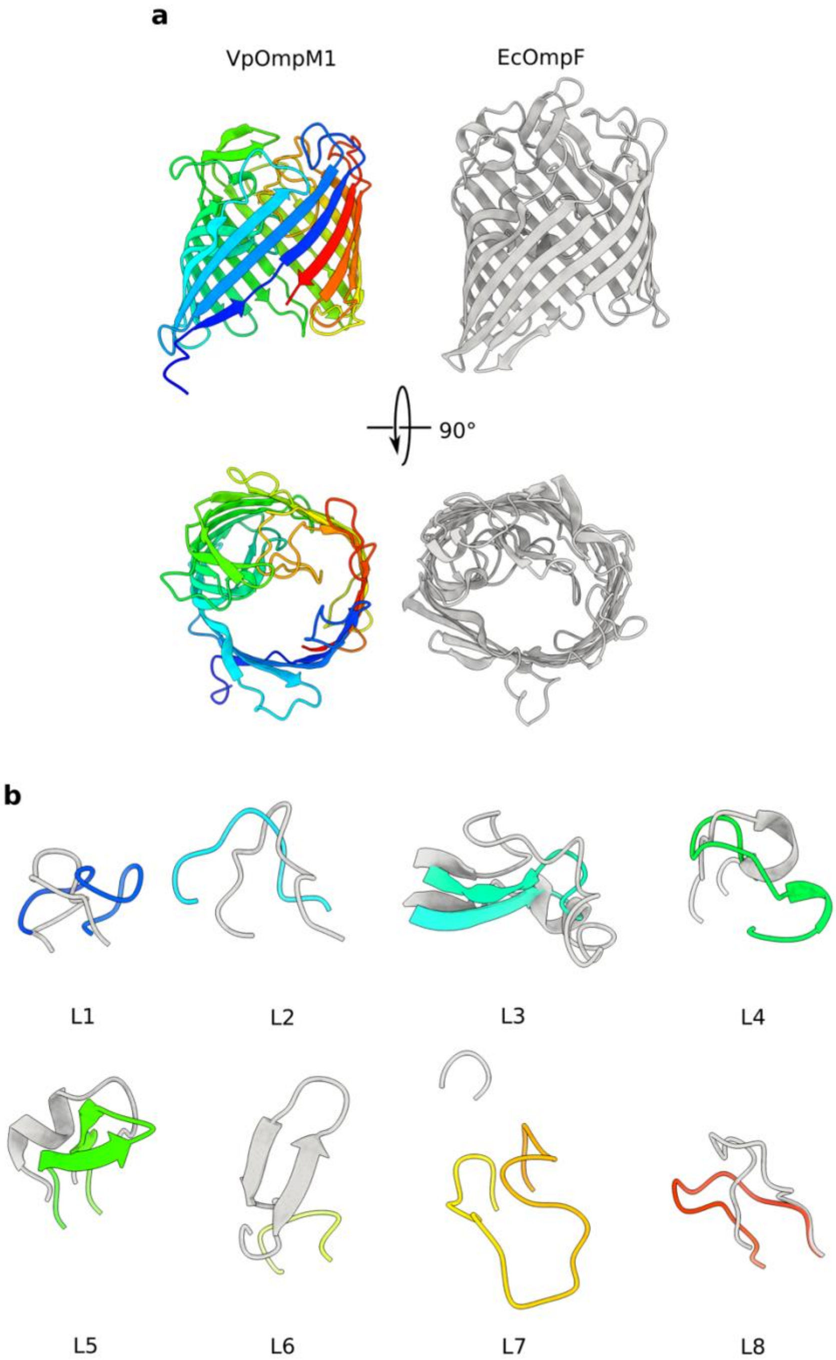


**Supplementary Figure 10. Comparison of VpOmpM1 and EcOmpF loops**. **a** VpOmpM1 β-barrel from C3 reconstruction (rainbow, N-terminus blue, C-terminus red) and EcOmpF (3POQ). Views generated from a superposition. **b** Comparison of the extracellular loops (L1-8) of the two β-barrels as viewed from inside the β-barrel.

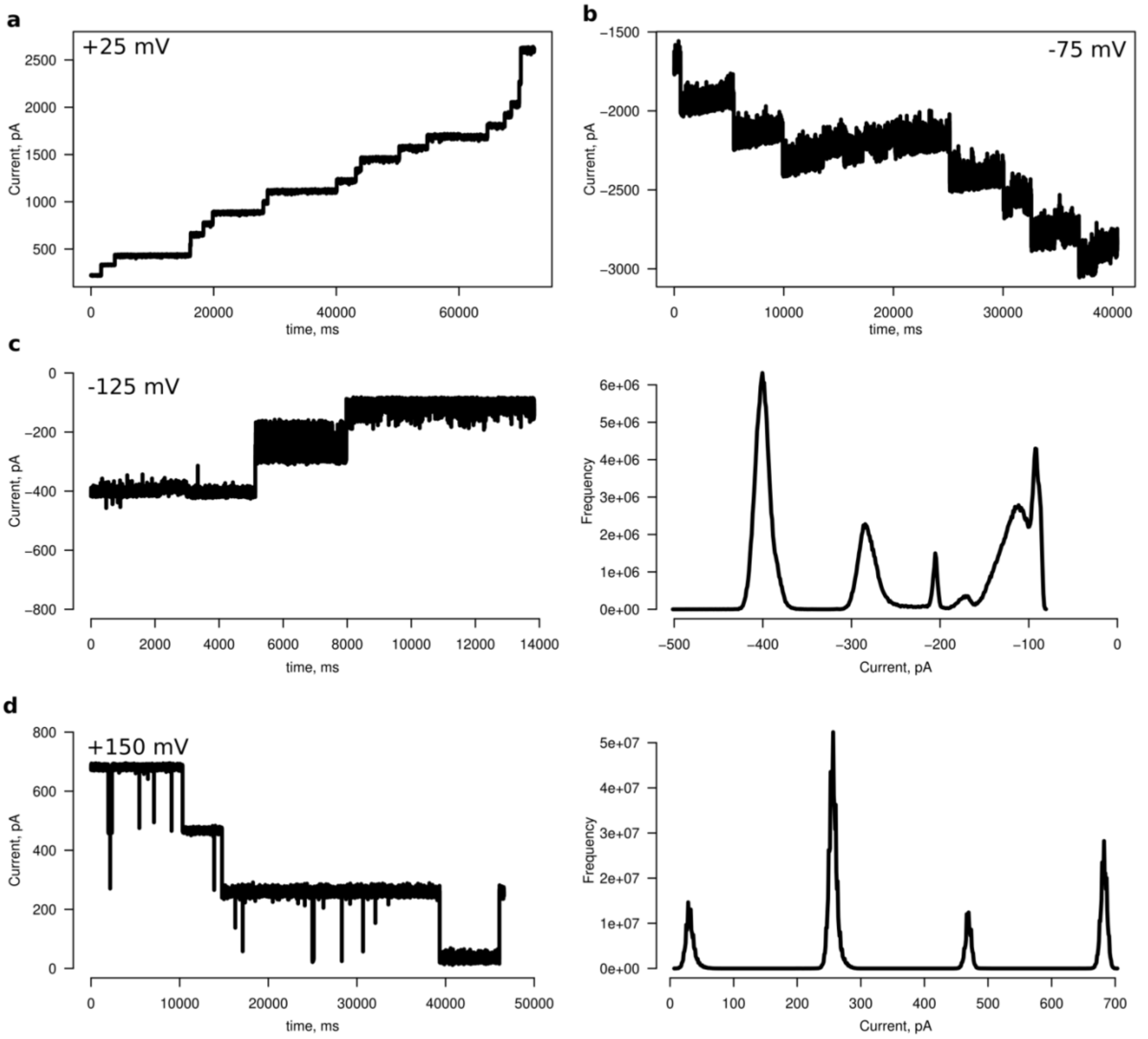


**Supplementary Figure 11. Representative electrophysiology recordings**. **a, b** Representative ion-current traces of full-length VpOmpM1 (**a**) and the barrel-only construct (**b**) showing multiple channel insertion events observed as ‘steps’ or sudden increases in current. **c, d** Ion-current traces for full-length VpOmpM1 (**c**) and the barrel-only construct (**d**) showing channel gating activity at high potential (left), with the corresponding all-current point histograms shown on the right. A 2.5 kHz low-pass eight-pole Bessel filter was applied to all traces. The applied voltage is indicated on each trace.

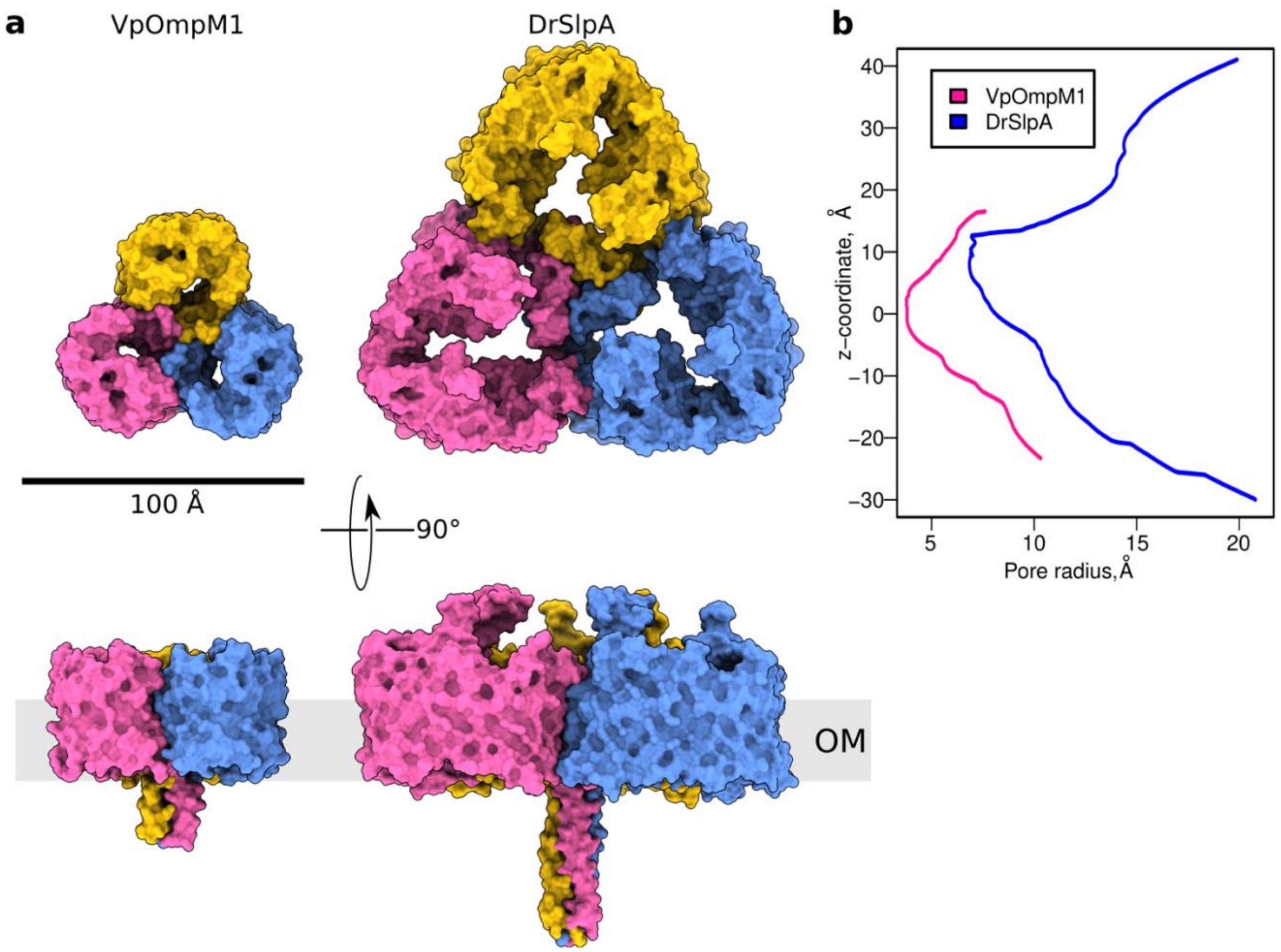


**Supplementary Figure 12. Comparison of VpOmpM1 and DrSlpA structures**. **a** To-scale surface representations of VpOmpM1 and *Deinococcus radiodurans* SlpA (DrSlpA) (PDB 8AE1) [3] from outside the cell (top) and from the OM plane (bottom). **b** HOLE [4] profiles of VpOmpM1 and DrSlpA. N.B. The profiles show the path of a ball-shaped probe via the narrowest part of the pore. Note that this analysis underestimates the size of the SlpA pore, since the constriction is not roughly circular as in VpOmpM1.

**Supplementary Movie 1.** All-atom MD simulation of the native VpOmpM1 structure with the grafted AlphaFold2-predicted stalk domain over 1 μs. Left, view parallel to the OM plane; right, view from the periplasm towards the OM. N.B. only the coiled-coil part of the stalk is shown.

**Supplementary Movie 2.** All-atom MD simulation of the compact structure predicted by AlphaFold2.

**Supplementary Movie 3.** A replicate of the all-atom MD simulation of the native VpOmpM1 structure with the grafted AlphaFold2-predicted stalk domain, in which the SLH domain of one protomer unfolds and contacts the coiled-coil.

**Supplementary Table 1.** Cryo-EM data collection, processing and refinement statistics.

| | **VpOmpM1 from *E. coli*** | | **VpOmpM1 from *V. parvula*** |
|---|---|---|---|
| **Data collection** | | | |
| Electron microscope | FEI Glacios | | FEI Glacios |
| Voltage (kV) | 200 | | 200 |
| Spherical aberration (µm) | 2.7 | | 2.7 |
| Camera | Falcon 4 (counting) | | Falcon 4 (counting) |
| Energy filter | none | | none |
| Magnification | 240,000 | | 240,000 |
| Pixel size (Å) | 0.574 | | 0.574 |
| Total dose ($e^-/Å^2$) | 50.1 | | 50 |
| Defocus minimum maximum (µm) | -1.0 to -2.0 | | -0.8 to -2.0 |
| Number of movies collected | 4,284 | | 6,505 |
| **Image Processing** | | | |
| Imposed symmetry | C1 | C3 | C1 |
| Initial number of particles | 838,051 | 838,051 | 1,501,693 |
| Final number of particles | 96,280 | 119,001 | 144,245 |
| Global resolution (FSC = 0.143) | 3.15 | 2.78 | 3.28 |
| Map sharpening B-factor ($Å^2$) | -70.2 | -85.5 | -97.5 |
| **Refinement** | | | |
| Model composition | | | |
| Non-hydrogen atoms | 7,386 | 7,077 | 7,406 |
| Protein residues | 963 | 924 | 968 |
| R.m.s. deviations | | | |
| Bonds lengths (Å) | 0.003 | 0.002 | 0.004 |
| Bond angles (˚) | 0.493 | 0.420 | 0.563 |
| Validation | | | |
| MolProbity score | 1.56 | 1.39 | 1.76 |
| Clash score | 4.52 | 3.57 | 6.45 |
| Rotamer outliers (%) | 0 | 0 | 0 |
| Ramachandran plot | | | |
| Favoured (%) | 95.30 | 96.41 | 93.97 |
| Outliers (%) | 0 | 0 | 0 |
| PDB | 8BYM | 8BYT | 8BYS |
| EMDB | EMD-16328 | EMD-16333 | EMD-16332 |

**Supplementary Table 2.** Crystallography data collection, processing and refinement parameters.

| | **VparOmpM1 stalk*** |
|---|---|
| **Data collection** | |
| DLS beamline | I03 |
| Wavelength | 0.89842 |
| Space Group | H3 |
| Unit cell parameters | |
| a, b, c (Å) | 118.71, 118.71, 47.24 |
| $\alpha$, $\beta$, $\gamma$ (°) | 90, 90, 120 |
| Molecules in AU | 3 |
| Resolution range (Å) | 34.78-1.57 (1.57-1.6) |
| I/$\sigma$I | 17.1 (0.7) |
| Completeness (%) | 99.5 (92.9) |
| Multiplicity | 9.0 (4.7) |
| Rpim (%) | 1.7 (88.4) |
| $CC_{1/2}$ (%) | 100 (23) |
| **Phasing** | |
| Ab initio | Arcimboldo[5] |
| **Refinement** | |
| Resolution (Å) | 34.78-1.7 |
| Rwork/Rfree (%) | 21.9 /24.8 |
| Reflections | |
| Non-hydrogen atoms | 2,015 |
| Protein only | 1,918 |
| Average B-factor (Å$^2$) | 39.68 |
| Rmsd | |
| Bond lengths (Å) | 0.011 |
| Bond angles (°) | 1.21 |
| MolProbity clashscore | 3.75 |
| Ramachandran plot | |
| Favoured (%) | 99.59 |
| Outliers (%) | 0 |
| PDB | 8BZ2 |

*Statistics for the highest-resolution shell are shown in parentheses.

**Supplementary Table 3.** Bacterial strains used in this study.

| Strain | Genotype | Reference or Source |
|---|---|---|
| ***E. coli* DH5α** | *F- endA1 glnV44 thi-1 recA1 relA1 gyrA96 deoR nupG purB20* φ80d*lacZ*ΔM15 Δ(*lacZYA-argF*)U169, hsdR17($r_K^- m_K^+$), $λ^-$ | Promega |
| ***E. coli* TOP10** | *F- mcrA Δ(mrr-hsdRMS-mcrBC) φ80lacZΔM15 ΔlacX74 nupG recA1 araD139 Δ(ara-leu)7697 galE15 galK16 rpsL(StrR) endA1 λ-* | Invitrogen |
| ***E. coli* BL21(DE3)** | *F- ompT lon hsdSB (rB- mB-) gal dcm* (DE3) | Invitrogen |
| ***E. coli* C43(DE3) *ΔcyoABCD*** | *F- ompT lon hsdSB (rB- mB-) gal dcm* (DE3) *lacUV5** ΔcyoABCD* | [6] (modified) |
| ***E. coli* MFDpir** | *MG1655 RP4-2-Tc ::[Δmu1 ::aac(3)IV-ΔaphA-Δnic35-Δmu2 ::zeo] ΔdapA :erm-pir) ΔrecA* | [7] |
| ***V. parvula* SKV38** | Wild type strain | [8] |
| ***V. parvula* SKV38 Δ*ompM1-3*** | *ΔFNLLGLLA_01389-7::tetM* | [9] |

**Supplementary Table 4.** Plasmids used in this study.

| Plasmid | Description | Reference or Source |
|---|---|---|
| **pB22** | pBAD22 vector with *E. coli* TamA signal sequence, N-terminal $His_7$-tag | [10,11] |
| **pJW45** | pB22 with beta barrel of *ompM1* cloned between XhoI and XbaI restriction sites | This study |
| **pJW46** | pB22 with *ompM1* (without signal peptide) CDS cloned between XhoI and XbaI restriction sites | This study |
| **pJW48** | pRPF185 with His-tagged *ompM1* (native RBS) cloned into the *Sac*I restriction site | This study |
| **pRPF185** | *Escherichia/Clostridium* shuttle conjugative expression vector with *tet* promoter | [12] |
| **pBAD24** | Vector for expressing proteins with their native signal sequence in *E. coli* | [10] |
| **pBAD24-EcOmpF** | pBAD24 vector with the full-length *E. coli* OmpF coding sequence cloned between EcoRI and XbaI restriction sites | This study |
| **pET28b** | *E. coli* expression vector | Novagen |
| **pET28b-SLH_22-107** | pET28b with *ompM1* SLH domain and stalk residues 22-107 coding region cloned between NcoI and XhoI sites (C-terminal $His_6$-tag) | This study |
| **pB22-00518_21-200** | pB22 with beta barrel (residues 21-200) of *FNLLGLLA_00528* cloned between XhoI and XbaI restriction sites | This study |
| **pB22-00833** | pB22 with beta barrel of *FNLLGLLA_00833* cloned between XhoI and XbaI restriction sites | This study |

**Supplementary Table 5**. Primers* used in this study.

| Primer | Sequence | Description |
|---|---|---|
| **JW172** | agcgttaacagatctgagcttaagaaagaaggaattcattatgaaaaaac | Forward primer for amplifying full length *ompM1* with its native RBS with homology arm to the pRPF185 digested with SacI |
| **JW201** | ccatcaccatcaccatcacctcgagcgcttggaagaccgtgtag | Forward primer for amplifying OmpM1 beta barrel with homology arm to the pB22 digested with XhoI |
| **JW202** | ctgaagcgggtgtgaatactctagattagaatttgtaatttaattctgcacg | Reverse primer for amplifying *ompM1* with homology arm to the pB22 digested with XbaI |
| **JW203** | ccatcaccatcaccatcacctcgaggctgcaaatccattctcc | Forward primer for amplifying *ompM1* without signal peptide with homology arm to the pB22 digested with XhoI |
| **JW206** | tctcctttactgcaggagct**ttaatgatgatgatgatgatgaccacc**gaatttgtaatttaattctgcacggtag | Reverse primer for amplifying *ompM1*, adding C-terminal 6His tag (indicated in bold) with homology arm to the pRPF185 digested with SacI. |
| **00833_F** | atactcgaggcatttgctgcagctcc | Forward primer for amplifying *FNLLGLLA_00833* without signal peptide with XhoI site |
| **00833_R** | atatctagattagaaggagtagcctacacc | Reverse primer for amplifying *FNLLGLLA_00833* without signal peptide with XbaI site |
| **00518_F** | atactcgagactcctcaaactcaattcaataaag | Forward primer for amplifying *FNLLGLLA_00518* barrel with XhoI site |
| **00518_R** | atatctagattaaccaccgaaacggtaagataatc | Reverse primer for amplifying *FNLLGLLA_00518* barrel with XbaI site |
| **stalk_F** | ataccatggctgcaaatccattctccg | Forward primer for amplifying *ompM1* stalk region with NcoI site |
| **stalk_R** | atactcgagctttacattacctacacggtc | Reverse primer for amplifying *ompM1* stalk region with XhoI site |

* The part of the primer hybridizing to the template is indicated by an underscore.

**Supplementary Table 6.** Position restraints, timesteps (dt), and durations of the equilibration phases in the all-atom MD simulations.

| Equilibration phase | Position restraint / kJ mol$^{-1}$ nm$^{-2}$ | | | | dt/ fs | Duration / ns |
|---|---|---|---|---|---|---|
| | Protein Backbone | Protein Sidechains | Lipid Headgroups | Dihedrals | | |
| **NVT1** | 4000 | 2000 | 1000 | 1000 | 1 | 0.125 |
| **NVT2** | 2000 | 1000 | 400 | 400 | 1 | 0.125 |
| **NPT1** | 1000 | 500 | 400 | 200 | 2 | 0.5 |
| **NPT2** | 500 | 200 | 200 | 200 | 2 | 0.5 |
| **NPT3** | 200 | 50 | 40 | 100 | 2 | 0.5 |
| **NPT4** | 50 | - | - | - | 2 | 0.5 |

**Supplementary Table 7.** Substrate concentrations used in liposome swelling assays.

| Substrate | Concentration (mM) |
|---|---|
| **Figure 5** – VpOmpM1, VpOmpM1 barrel and EcOmpF | |
| **Lactate** | 8 |
| **Acetate** | 8 |
| **Putrescine** | 8 |
| **Arginine** | 8 |
| **Lysine** | 8 |
| **Glutamate** | 10 |
| **Aspartate** | 8 |
| **Glycine** | 15 |
| **Alanine** | 15 |
| **Leucine** | 12 |
| **Methionine** | 15 |
| **Arabinose** | 12 |
| **Glucose** | 12 |
| **Fructose** | 12 |
| **Lactose** | 15 |
| **Maltose** | 15 |
| **Sucrose** | 15 |
| **Kanamycin** | 8 |
| **Ampicillin** | 8 |
| **Gentamicin** | 8 |
| **Figure 6** – FNLLGLLA_00518 barrel, VpOmpM1 barrel and EcOmpF | |
| **Lactate** | 10 |
| **Putrescine** | 8 |
| **Glycine** | 15 |
| **Arabinose** | 10 |

**Supplementary references**


1. Holm, L. Dali server: structural unification of protein families. *Nucleic Acids Res.* **50**, W210–W215 (2022).
2. Jumper, J. *et al.* Highly accurate protein structure prediction with AlphaFold. *Nat. 2021 5967873* **596**, 583–589 (2021).
3. von Kügelgen, A., van Dorst, S., Alva, V. & Bharat, T. A. M. A multidomain connector links the outer membrane and cell wall in phylogenetically deep-branching bacteria. *Proc. Natl. Acad. Sci. U. S. A.* **119**, e2203156119 (2022).
4. Smart, O. S., Neduvelil, J. G., Wang, X., Wallace, B. A. & Sansom, M. S. P. HOLE: A program for the analysis of the pore dimensions of ion channel structural models. *J. Mol. Graph.* **14**, 354–360 (1996).
5. Rodríguez, D. D. *et al.* Crystallographic ab initio protein structure solution below atomic resolution. *Nat. Methods 2009 69* **6**, 651–653 (2009).
6. Miroux, B. & Walker, J. E. Over-production of Proteins inEscherichia coli: Mutant Hosts that Allow Synthesis of some Membrane Proteins and Globular Proteins at High Levels. *J. Mol. Biol.* **260**, 289–298 (1996).
7. Ferrières, L. *et al.* Silent mischief: bacteriophage Mu insertions contaminate products of Escherichia coli random mutagenesis performed using suicidal transposon delivery plasmids mobilized by broad-host-range RP4 conjugative machinery. *J. Bacteriol.* **192**, 6418–6427 (2010).
8. Knapp, S. *et al.* Natural Competence Is Common among Clinical Isolates of Veillonella parvula and Is Useful for Genetic Manipulation of This Key Member of the Oral Microbiome. *Front. Cell. Infect. Microbiol.* **7**, (2017).
9. Witwinowski, J. *et al.* An ancient divide in outer membrane tethering systems in bacteria suggests a mechanism for the diderm-to-monoderm transition. *Nat. Microbiol.* **7**, 411–422 (2022).
10. Guzman, L. M., Belin, D., Carson, M. J. & Beckwith, J. Tight regulation, modulation, and high-level expression by vectors containing the arabinose PBAD promoter. *J. Bacteriol.* **177**, 4121–4130 (1995).
11. Van den Berg, B. *et al.* X-ray structure of a protein-conducting channel. *Nature* **427**, 36–44 (2004).
12. Fagan, R. P. & Fairweather, N. F. Clostridium difficile Has Two Parallel and Essential Sec Secretion Systems. *J. Biol. Chem.* **286**, 27483–27493 (2011).
